# Assembly and analysis of the genome of *Notholithocarpus densiflorus*

**DOI:** 10.1101/2023.12.20.572644

**Authors:** Ying Cai, Ellis Anderson, Wen Xue, Sylvia Wong, Luman Cui, Xiaofang Cheng, Ou Wang, Qing Mao, Sophie Jia Liu, John T. Davis, Paulo R. Magalang, Douglas Schmidt, Takao Kasuga, Matteo Garbelotto, Radoje Drmanac, Chai-Shian Kua, Charles Cannon, Julin N. Maloof, Brock A. Peters

**Affiliations:** Advanced Genomics Technology Laboratory, Complete Genomics, 2904 Orchard Parkway, San Jose, CA 95134; BGI-Shenzhen, Shenzhen 518083, China; MGI, BGI-Shenzhen, Shenzhen 518083, China; Department of Plant Biology, University of California, One Shields Avenue, Davis, CA, 95616, USA; Department of Environmental Science, Policy & Management, University of California, Berkeley, California, USA; Crops Pathology & Genetics Research Unit, United States Department of Agriculture–Agricultural Research Service, Davis, CA 95616; Center for Tree Science, The Morton Arboretum, 4100 Illinois Route 53, Lisle, IL 60532

## Abstract

Tanoak (*Notholithocarpus densiflorus*) is an evergreen tree in the Fagaceae family found in California and southern Oregon. Historically, tanoak acorns were an important food source for Native American tribes and the bark was used extensively in the leather tanning process. Long considered a disjunct relictual element of the Asian stone oaks (*Lithocarpus spp*.), phylogenetic analysis has determined that the tanoak is an example of convergent evolution. Tanoaks are deeply divergent from oaks (*Quercus*) of the Pacific Northwest and comprise a new genus with a single species. These trees are highly susceptible to ‘sudden oak death’ (SOD), a plant pathogen (*Phytophthora ramorum*) that has caused widespread mortality of tanoaks. Here, we set out to assemble the genome and perform comparative studies among a number of individuals that demonstrated varying levels of susceptibility to SOD. First, we sequenced and *de novo* assembled a draft reference genome of *N. densiflorus* using co-barcoded library processing methods and an MGI DNBSEQ-G400 sequencer. To increase the contiguity of the final assembly, we also sequenced Oxford Nanopore (ONT) long reads to 30X coverage. To our knowledge, the draft genome reported here is one of the more contiguous and complete genomes of a tree species published until now, with a contig N50 of ∼1.2 Mb and a scaffold N50 of ∼2.1 Mb. In addition, we sequenced 11 genetically distinct individuals and mapped these onto the draft reference genome enabling the discovery of almost 25 million single nucleotide polymorphisms and ∼4.4 million small insertions and deletions. Finally, using co-barcoded data we were able to generate complete haplotype coverage of all 11 genomes.

## Introduction

Tanoak (*Notholithocarpus densiflorus* (Manos et al. 2008) is a compelling part of the beech family (Fagaceae) and possesses an unusual evolutionary history. Long considered a disjunct relictual element of the Asian stone oaks (*Lithocarpus*), modern phylogenetic analysis determined that the tanoak was a clear example of convergent evolution in fruit type, requiring the recognition of a new genus comprising a single species (Manos et al. 2008). More recent phylogenomic analyses (Zhou et al. 2022) places it basal and sister to all northern hemisphere oaks (genus *Quercus*), both Old and New World groups. Tanoak is also the last common ancestor with insect pollination in a species-rich wind-pollinated clade, splitting with the oaks roughly 54 million years ago. Little fossil evidence for the taxon exists but its current geographic distribution is restricted to a relatively small area in the Pacific Northwest, suggesting that taxon may belong to been a species-poor clade for a significant period of time. By comparison, the North American oaks have diversified and spread throughout North America, occupying a wide range of habitats (Hipp et al. 2018).

Ecologically, tanoaks are adapted to a Mediterranean-type climate, with a long dry season and periodic fires. They can tolerate a wide range of soil types, from shallow rocky soils to deep, well-drained soils. Two growth forms exist, recognized as different varieties: *Notholithocarpus densiflorus* var. *densiflorus* is a tree, with individuals growing to 45 m in height, often as a co-dominant in the redwood and mixed evergreen forests of the north coast ranges, while *Notholithocarpus densiflorus* var. *echinoides* is a shrub, more commonly growing at higher elevations in open conifer forests and dry slopes of the northern interior. As a locally dominant species in these habitats, *Notholithocarpus* trees play an important ecosystem role, forming the mid- and lower canopy strata of redwood forests and providing habitat and food for a variety of wildlife, including birds and mammals (Waring and O’Hara 2008). Additionally, their thick bark and ability to resprout from the base after fire or other damage make them an important component of the forest’s resilience and recovery.

Tanoak obtained its common name from the extensive harvest of their bark during the early 20^th^ century for the regional tanning industry (Bowcutt 2011), a business that paradoxically first led to increased tanoak densities due to prolific coppicing, and then made it a frequent target of herbicide applications to reduce densities. Both the acorns and the bark of Notholithocarpus trees have been used for food and leather processing by indigenous peoples in North America for centuries (Bowcutt 2015). SOD, caused by the oomycete pathogen, *Phytophthora ramorum*, has killed tens of millions of tanoak, coast live oak, California black oak, and other native tree species (Aphis.usda.gov). Tanoak is the most susceptible species to *P. Ramorum* (Davidson et al. 2003). Its decline due to the rapid spread of SOD has the potential to dramatically affect the overall biodiversity and conservation status of these forests (Cobb et al. 2012), particularly compromising their role as one of the few local ectomycorrhizal hosts (Bergemann and Garbelotto 2006). Overall, the loss of tanoak from redwood forests will reduce biodiversity and alter fundamental ecosystem processes (McCallum and Dobson 1995; Rizzo et al. 2005; Wardle et al. 2011).

The difference in the susceptibility to SOD between Notholithocarpus and Quercus is also a compelling question, potentially associated with the dramatic differences in their evolutionary history and reproductive biology. North American oak species generally participate in a large continental scale syngameon (Cannon et al. In review), which potentially enhances the overall diversity found in their genome and particularly in disease resistance genes (Cannon and Petit 2020). This genetic exchange among oak species is probably facilitated by their wind-pollination, in comparison to the insect-pollinated tanoak. The existence of a single species of tanoak obviously prevents it from gaining any evolutionary advantage from participation in a syngameon, no matter its pollination syndrome. This substantial difference between these two approximately similar-aged lineages – the species rich syngameon of the oaks versus the species poor (monospecific?) isolate of the tanoak – should have considerable impact on the overall genomic evolution and potential susceptibility to SOD. We set out to assemble the genome of tanoak and perform comparative studies among a number of individuals that demonstrated varying levels of susceptibility to SOD. We then compared this assembly to existing completed genomes in the *Fagaceae*. The questions we asked were:

1. Do basic genomic properties differ between species-rich and species-poor lineages?
2. Can differences in the overall diversity of disease resistance genes be detected?

## Results

Thirteen individual trees, 11 of which are genetically distinct and from disparate locations, were selected in order to help gauge the diversity within th**e** *N. densiflorus* species (Figure 1a). In addition, trees were selected based on various levels of susceptibility to *Phytophthora ramorum*, the plant pathogen that causes Sudden Oak Death, ranging from relatively resistant to highly susceptible (Supplementary Table 1). Genomic DNA from a leaf of each tree was used to make co-barcoded sequencing libraries using the single-tube long fragment read (stLFR) process (Wang et al. 2019; Xiaofang et al. 2018). Approximately 100 Gb of data per sample were generated using an MGI DNBSEQ-G400 second generation DNA sequencer (Supplementary Table 1). Reads from each sample were analyzed with GenomScope (Vurture et al. 2017) to determine the kmer spectra and heterogeneity of each sample as well as estimate the size of the *N. densiflorus* genome (Supplementary Table 2 and Supp Figure 1). Both the estimated size and kmer heterogeneity fell within the range of other closely related species (Supplementary Table 3). The genomic DNA of six samples was further enriched for higher molecular weight fragments (labeled with XL) and additional stLFR libraries were made and processed as above but with ∼200 Gb of data generated per sample. These were individually *de novo* diploid assembled using a modified version of 10X Genomics’ Supernova software (Weisenfeld et al. 2017) resulting in contig and scaffold N50 values ranging from 31.4-50.4 kb and 0.145-2.05 Mb, respectively (Supplementary Table 4). Using Merqury (Rhie et al. 2020), a kmer based assembly analysis program, a per base quality score ranging from Q50-59 and an estimated completeness of 82-89% (Supplementary Table 4) were generated for each of the six samples. A single pseudo haplotype with the overall best assembly (contig N50 of 50.4 kb, scaffold N50 of 2.05 Mb, 89% complete, and Q59) was selected (SM.74.45.XL (clone 1)) for use as the draft reference *N. densiflorus* genome and contigs from the remaining five XL assemblies were used to fill gaps within each scaffold of the draft using TGS-GapCloser (Xu et al. 2020). This resulted in a large improvement in contiguity from an N50 of 50.4 kb to 385.8 kb (Table 1). To further increase contiguity, 27 Gb of nanopore data (ONT) from NL2.XL, SM.52.81.XL (clone 2), and SM.54.37XL were used to fill remaining gaps and achieve a contig N50 of ∼1 Mb (Table 1). This assembly was further polished to remove errors using Pilon (Walker et al. 2014) with the NGS read set from SM.74.45.XL (clone 1). Purge Haplotigs (Roach et al. 2018) was used to remove duplicated regions in the genome and resulted in a reduction in size from 916.6 Mb to 777.5 Mb. This is closer to the expected size of 785 Mb as determined by flow cytometry (Supplementary Figure 2). In addition, the entire set of ONT reads was assembled using Flye v2.9 (Kolmogorov et al. 2019). This assembly was aligned with the draft reference in order to find potential insertions and deletion errors (Supplementary Figure 3). Overall, the two assemblies aligned very closely with only one large region found to be duplicated in the draft reference versus the ONT assembly. Further inspection of read coverage in this region, after mapping all of the samples to the draft reference, suggested that this duplication is present in the tanoak genome and should not be removed. In each case raw ONT reads were used to determine what corrective actions should be taken. Finally, BUSCO (Manni et al. 2021) analysis was performed on the draft genome resulting in a complete gene score of 95.5% with a low duplication rate of 4.5% (Table 1). In addition, dot plots between the tanoak draft genome and two related species of Oak (*Q. robur* and *Q. rubra*, Figure 1b and 1c) showed close alignment. Taken together, these results suggest that we have generated a high-quality draft reference genome for *N. densiflorus*.

**Table 1.**
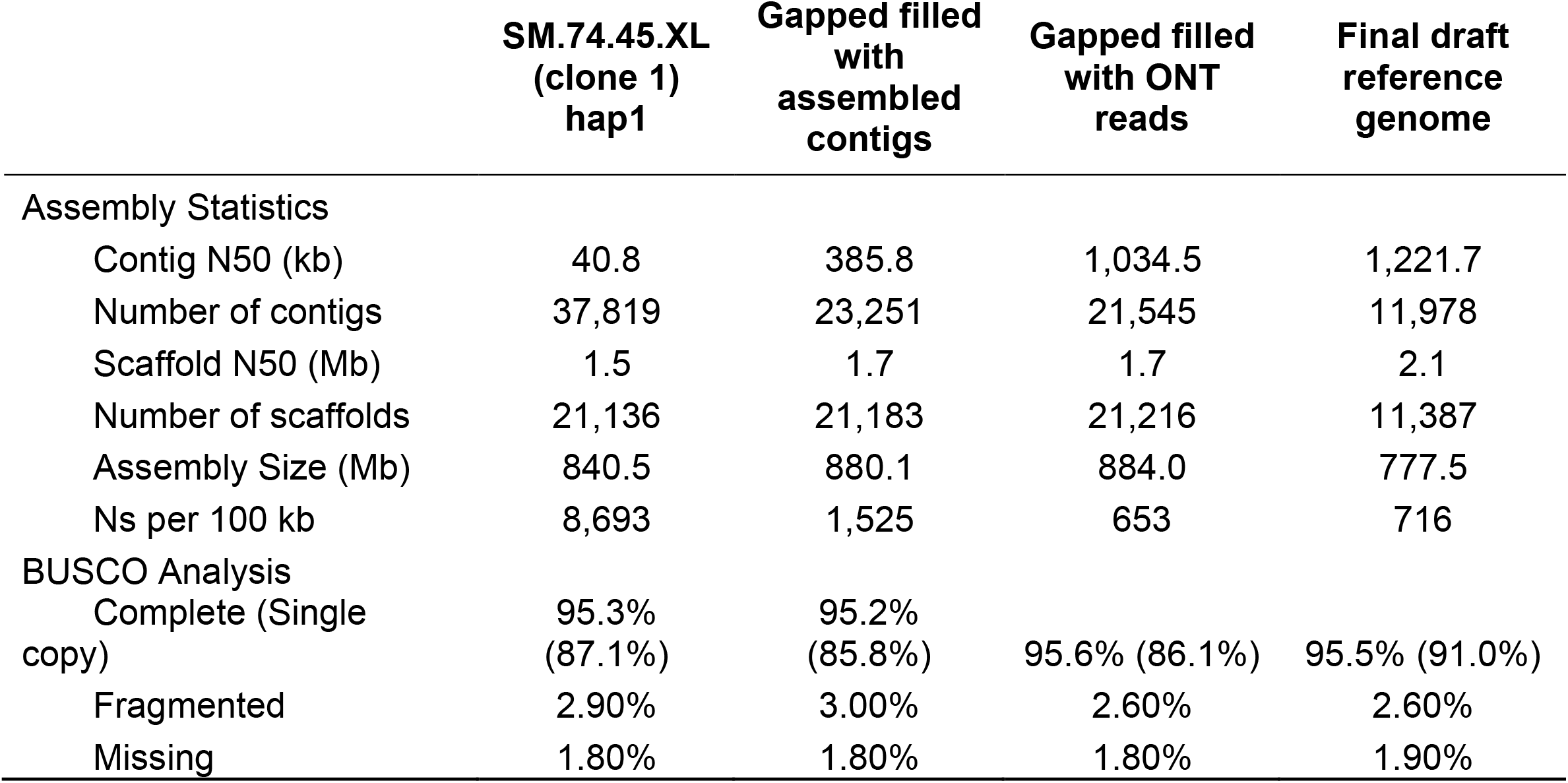
Assembly statistics.

**Table 2.**
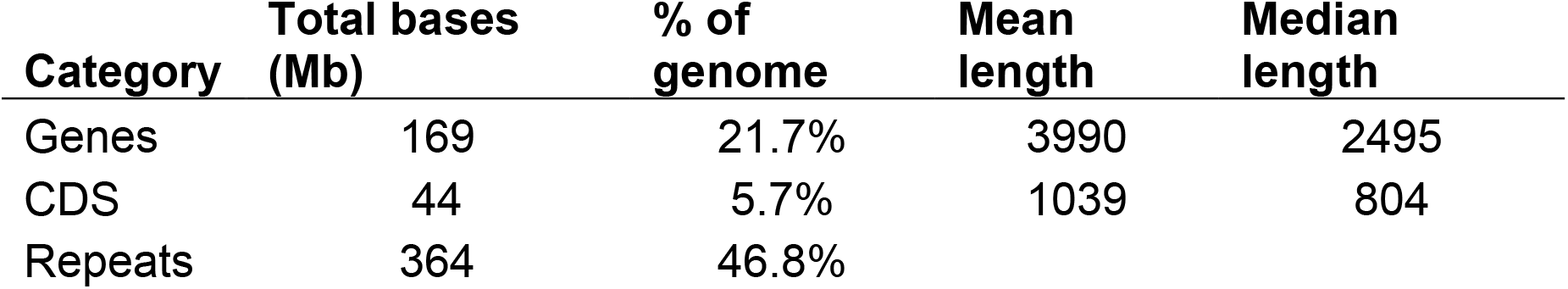
Annotation statistics.

**Table 3.** Tanoak R gene content versus other species.

| Species | TNL<sup>1</sup> | CNL<sup>2</sup> | Total | Complete | Complete pseudo | Partial | Partial pseudo | Genome Size (Mb) | Complete /Genome Size |
| --- | --- | --- | --- | --- | --- | --- | --- | --- | --- |
| Walnut (*Juglans regia*) | 199 | 145 | 460 | 251 | 129 | 27 | 53 | 573 | 0.44 |
| Chinese Chestnut (*Castanea mollissima*) | 143 | 188 | 418 | 242 | 115 | 31 | 30 | 413 | 0.59 |
| **Tanoak (*Notholithocarpus densiflorus*)** | **302** | **398** | **947** | **505** | **248** | **126** | **68** | **778** | 0.65 |
| Poplar (*Populus simonii*) | 213 | 204 | 637 | 359 | 143 | 77 | 58 | 434 | 0.83 |
| Northern red oak (*Quercus rubra*) | 354 | 510 | 1320 | 613 | 327 | 77 | 88 | 740 | 0.83 |
| Grape (*Vitis vinifera*) | 174 | 336 | 739 | 416 | 202 | 74 | 47 | 486 | 0.86 |
| Pendunculate Oak (*Quercus robur*) | 494 | 554 | 1319 | 773 | 360 | 102 | 84 | 814 | 0.95 |
| *Arabidopsis thaliana* | 115 | 30 | 171 | 122 | 20 | 21 | 8 | 120 | 1.02 |
| Peach (*Prunus persica*) | 166 | 186 | 415 | 257 | 92 | 37 | 29 | 227 | 1.13 |
| European beech (*Fagus sylvatica*) | 376 | 547 | 1290 | 699 | 375 | 112 | 104 | 541 | 1.29 |

<sup>1</sup>TIR-domain-containing
<sup>2</sup>CC-domain-containing

**Figure 1.**
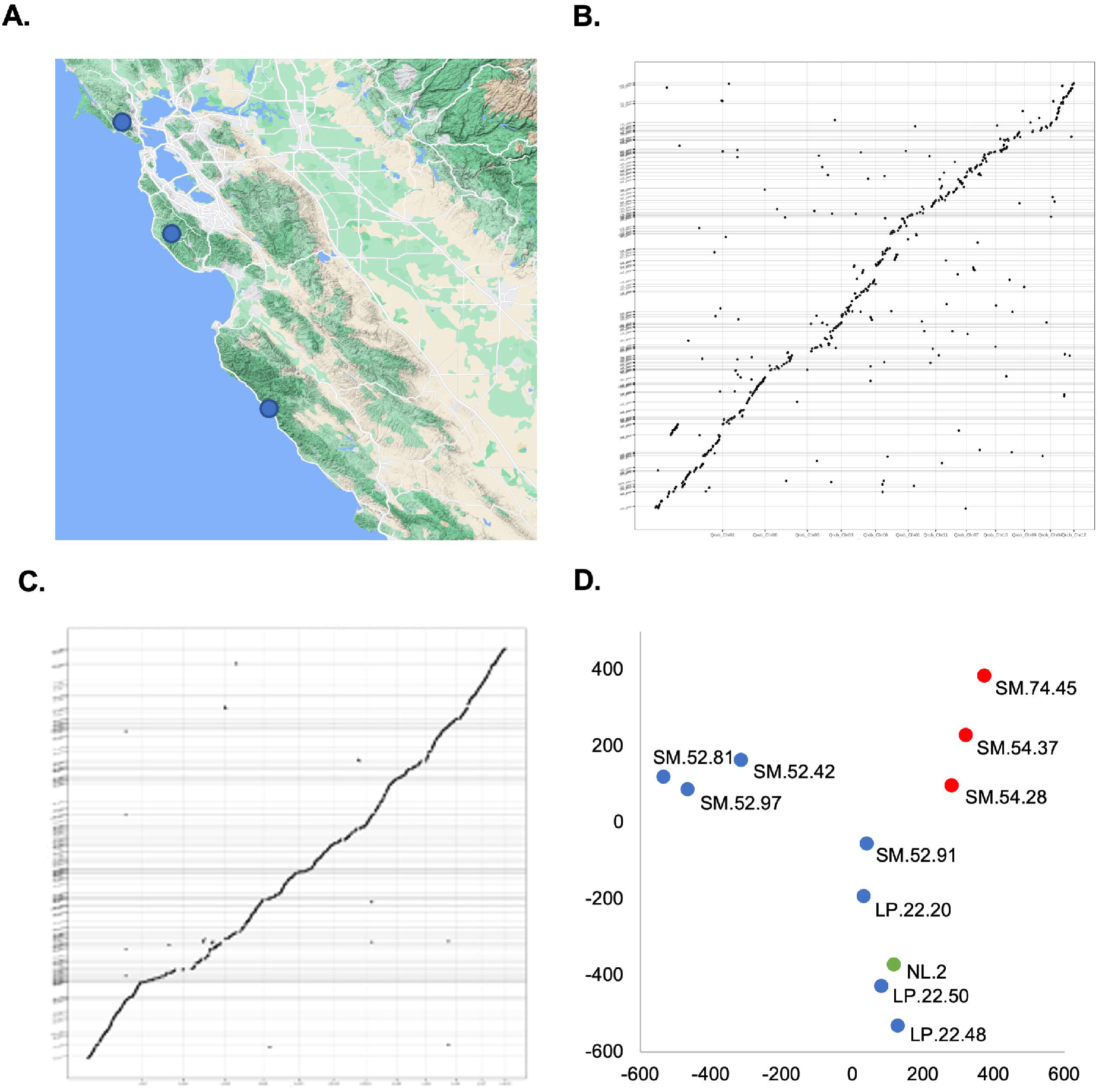
*N. densiflorus* project characteristics. **A**. Samples were collected in multiple locations across Central California as displayed on the map. Assembled contigs from the draft Tanoak reference (y-axis) were compared against the *Q. robur* (**B**) and *Q. rubra* (**C**) assemblies (x-axis). The 11 genetically distinct tanoak samples were projected onto a PCA generated from 2.4 million bi-allelic SNPs (**D**). The first (x-axis) and second (y-axis) principal components were plotted for each sample. Tree specimens that have shown increased susceptibility to Phytophthora ramorum are colored red, those with increased resistance are colored blue, and those with unknown susceptibility are colored green.

**Figure 2.**
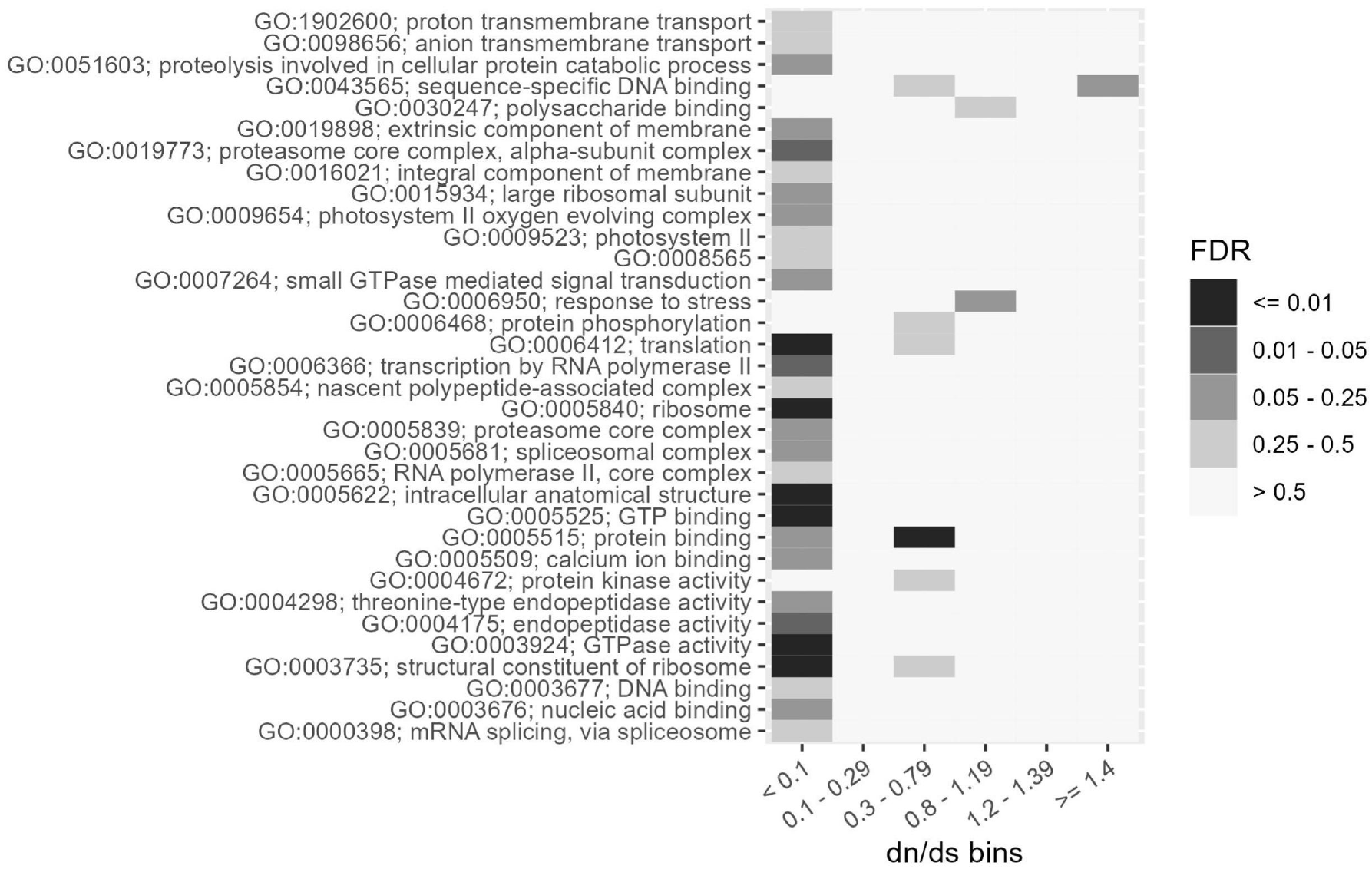
dN/dS analysis. Genes were binned based on their dN/dS ratio (x-axis) and then GO enrichment was performed for genes in each bin. Shading indicates the false discovery rate for GO enrichment.

**Figure 3.**
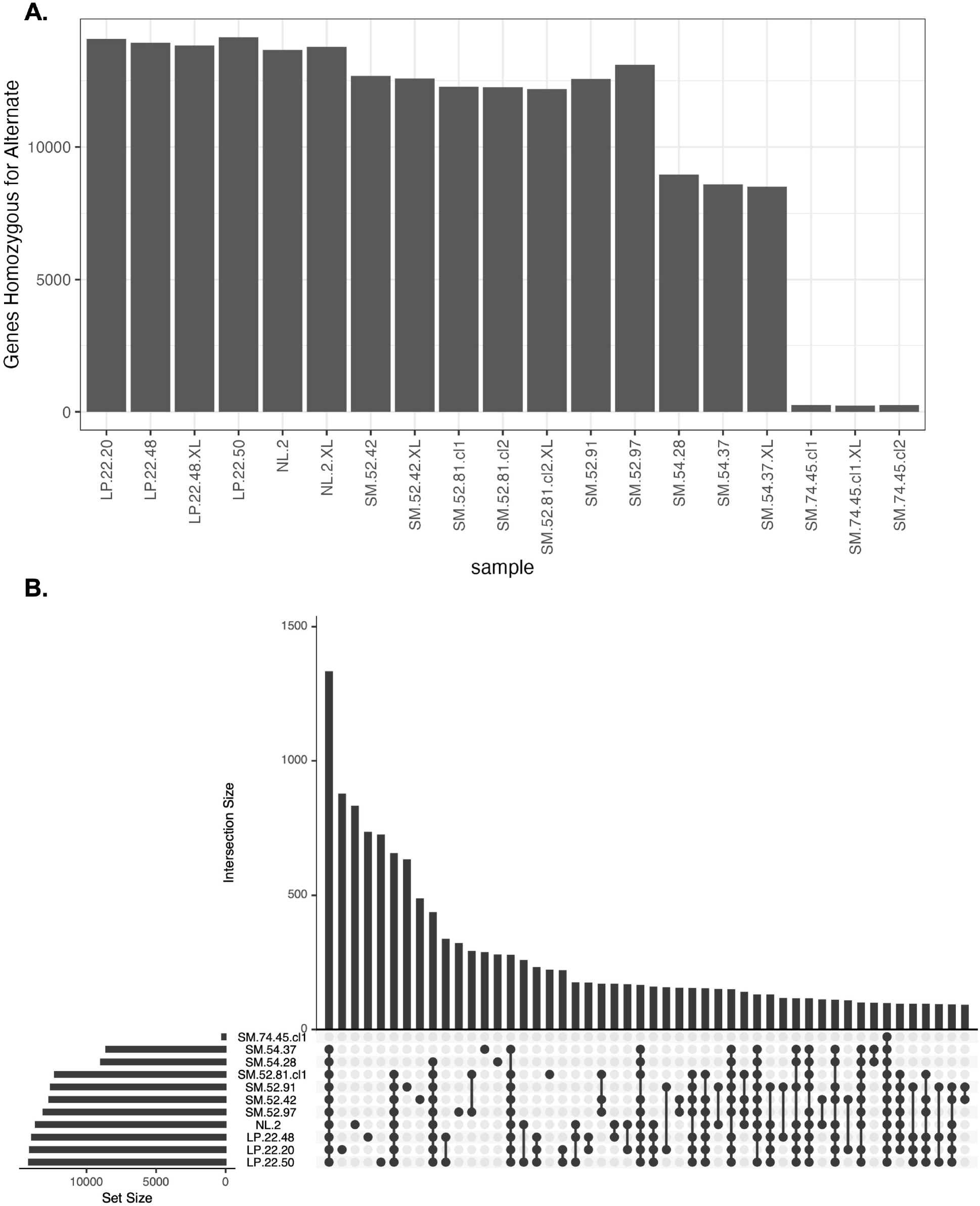
Distribution of moderate and high effect alleles. **A)** The number of genes with at least one moderate or high effect alternate allele (compared to SM.74.45 reference) for each sample. **B)** An UpSet plot (Conway et al. 2017; Lex et al. 2014) showing the unique and shared alleles among samples.

Next, we proceeded to annotate the coding sequences of the draft genome with MAKER (Campbell et al. 2014b), AUGUSTUS (Stanke and Waack 2003), SNAP (Korf 2004), protein sequence from the taxonomically close *Quercus robur* (Plomion et al. 2018), and a Trinity (Grabherr et al. 2011) *de novo* assembled transcriptome from a study of RNA sequencing data from 45 different *N. densiflorus* samples (Kasuga et al. 2021). This resulted in the placement of 42,319 genes onto the draft genome (Table 2). To explore the resistance (R) gene content (nucleotide-binding site leucine-rich repeat (NLR) genes), we used NLR-annotator (Steuernagel et al. 2020) and compared the tanoak results to the genomes of nine different plant species (Table 3). *Q. robur* has been found to be more resistant to *P. ramorum* and was found to have approximately 1.6-fold more complete non-pseudo R genes than *N. densiflorus* (Table 3). While intriguing, further studies will be necessary to determine the cause of increased resistance in *Q. robur*.

To look for genes that may have been subject to selection since the divergence of oak and tanoak, we calculated the ratio of nonsynonymous to synonymous substitutions(dN/dS) for all annotated tanoak genes for which we could identify a clear oak ortholog by reciprocal blast. A total of 5,541 were found to have a clear ortholog, GO annotation, and sequence variation between these species. To ask if particular types of genes were enriched in genes showing signs of purifying or positive selection, we binned genes based on their dN/dS value and calculated Gene Ontology (GO) enrichment for each bin (Figure 2). Three hundred and fifty-nine genes showed evidence of strong purifying selection (dN/dS < 0.1) in processes such as protein translation, ribosomes, protein degradation, and RNA pol II transcription, as expected based on the fundamental nature of these processes. With regards to positive selection, 201 genes were found to have a dN/dS ratio above 1.2; and the GO term “sequence-specific DNA binding” was marginally enriched for genes with a dN/dS >= 1.4 (FDR = 0.20). Interestingly, eight of the nine genes in this category had homology to *Arabidopsis* genes related to pathogen defense or abiotic stress (the ninth gene did not have a functionally annotated homolog; Supp Table 5). Specifically, there were three genes with homology to *Arabidopsis* WRKY transcription factors each implicated in microbial defense (WRKY 11, 40, and 41), three genes with homology to *Arabidopsis* genes regulated by abscisic acid (RAS1 and two ATHB7 homologs) and two genes with homology to *Arabidopsis* heat-stress transcription factors (AtHSFA-2 and 3).

In order to investigate the variation between samples we mapped all of the individual libraries onto the newly created reference genome. This resulted in the discovery of ∼25 million SNPs and of ∼4.4 million small insertions and deletions with an average of 7.7 million per individual tree. Principal component analysis (PCA) was performed using SNP data from all libraries (Figure 1d and Supplementary Figure 4). As expected, replicate libraries made from the same DNA sample and libraries from related family members tended to cluster with each other (Supplementary Figure 4). Projecting SOD susceptibility on the PCA did not inform beyond what was already known based on family inheritance (Figure 1d).

**Figure 4.**
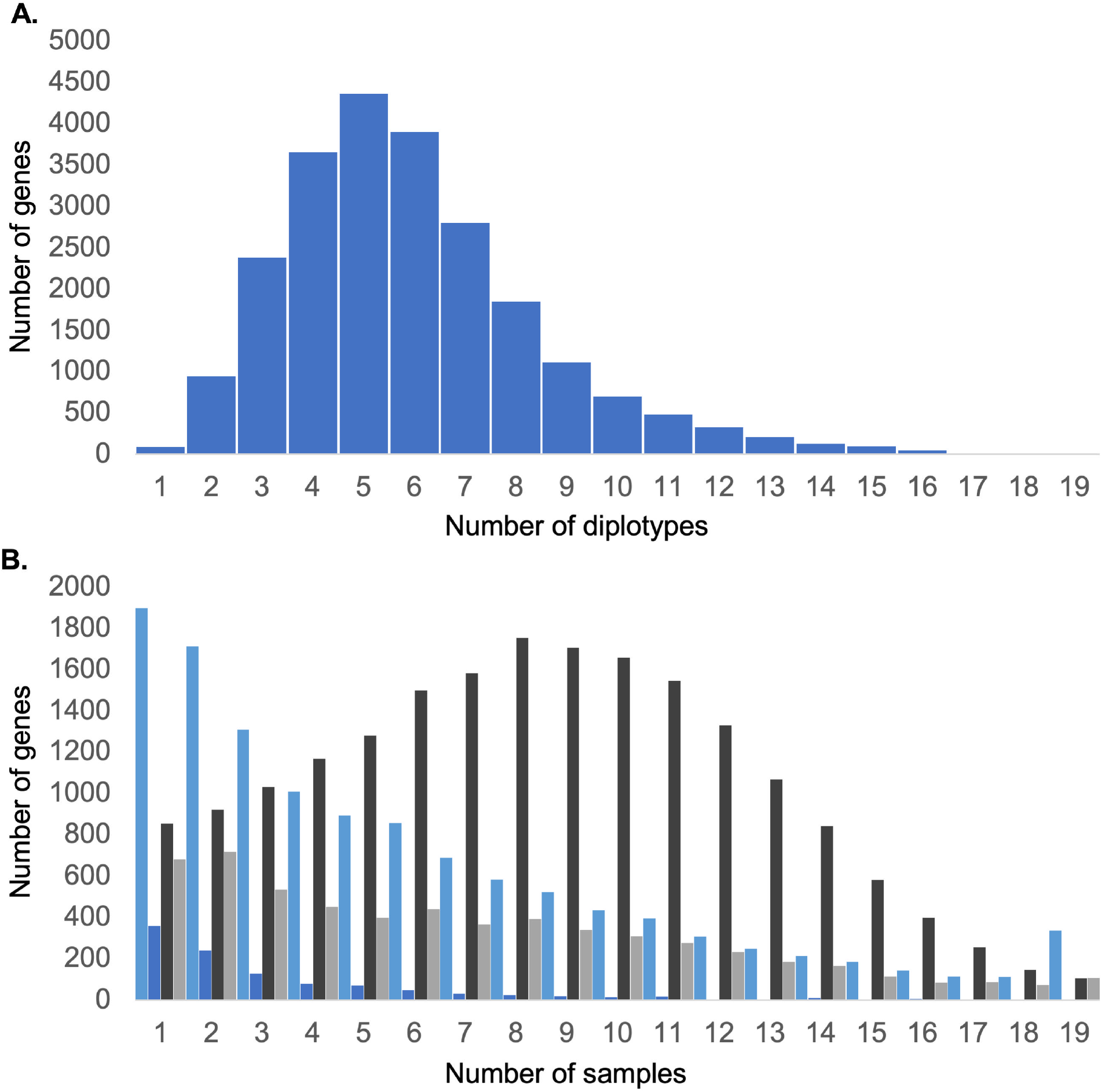
Diplotype analysis. The total number of diplotypes across the entire set of samples were calculated using haplotype information from each sample. **A)** Diplotypes per gene were calculated and summed resulting in a median of 6. **B)** The number of samples with at least one moderate (light blue) or high (blue) SnpEff called variant in each allele of a specific gene were summed. The same calculations were done for two or more moderate (dark grey) or high (grey) SnpEff called variants in the same allele of a specific gene.

Of the total variants identified in tanoak, 604,032 resulted in coding changes to 39,574 different genes. Using SnpEff (Cingolani et al. 2012), these were further evaluated resulting in the categorization of 526,584 SNPs as predicted to have a moderate impact on 38,837 genes and 787,687 high impact variants in 22,361 genes. Comparing the reference tree SM.74.45 to the other trees, we found that SM.54.28 and SM.54.37 differ from the reference at high and/or moderate alleles in 8,499 to 8,965 genes, and the remaining trees differed from the reference in 13,095 to 14,069 genes (Figure 3a). We also compared overlap of alternate alleles using an UpSet plot (Conway et al. 2017; Lex et al. 2014) (Figure 3b). Unsurprisingly, the most common category are those SNPs that are alternate in all trees except for the reference. Interestingly, the next four most common categories are SNPs that are unique to individual trees (or trees and clones), indicating a high degree of diversity among these trees. This plot reveals that each tree has a large number of unique SNPs. Additionally, when all libraries are added to this plot (Supplementary Figure 5) the concordance across clones and “XL” samples of the same tree are shown.

In an effort to identify any potential correlations to SOD resistance we compared genes with detrimental variants to susceptibility scores to *Phytophthora ramorum*. A total of 213 genes were found where the average nominal p-value for SNP association with disease susceptibility was < 0.05 (Fisher’s exact test), however given the relatively small sample size of our study these correlations were not statistically significant after correction for multiple testing. Nevertheless, there were several potentially interesting genes among these 213, including a homolog of *LIK1*, a leucine-rich-repeat receptor-like kinase (LRR-RLK) gene involved in plant innate immunity to microbes, a homolog of *COI1*, an essential component of jasmonic acid signaling, and *MYC2*, a key defense regulator. Further studies with a larger number of individual trees will be necessary to determine if any of these genes are associated with SOD resistance.

One of the unique advantages of using stLFR to analyze these samples is that genome wide haplotype data could be generated for all samples. Using HapCut2, an average haplotype contig N50 value of ∼1.6 Mb was achieved enabling the exploration of haplotype variation across the different samples (Supplementary Figure 6 and Supp Table 1) and enabling the determination of 136,541 diplotypes (a specific combination of two individual haplotypes) found across 23,089 genes, with a median of 6 diplotypes per gene (Figure 4a). Combining this information with the SnpEff analysis allowed the discovery of 188 genes on average per tree with moderate or high impact changes predicted in both alleles (Figure 4b). We further asked if there were any genes where susceptible trees had the reference allele and resistant trees were all heterozygous for two different moderate/high effect alleles (or vice versa). We found one gene with homology to the Arabidopsis gene AT1G59780, annotated as a putative disease-resistance protein, that had the reference sequence in all susceptible trees and was heterozygous for two different moderate or high impact changes in all resistant trees. We also found one gene with the opposite pattern; this gene had homology to AT1G08730, coding for a class XI myosin homolog. For both of these genes the association with disease resistance was nominally significant (Fisher’s exact test p-value = 0.002) but not significant after correcting for multiple testing.

## Discussion

In this study, we sequenced and *de novo* assembled a draft reference genome for the species *Notholithocarpus densiflorus*, a member of the beech family, using cobarcoded second generation reads. Using kmer analysis we estimate the initial assembly has approximately one error in 850,000 bases (Q59.3). We further refined and filled gaps in this assembly by adding contigs from other assembled tanoak samples as well as through the use of third generation continuous reads. The draft reference we present here is one of the most contiguous tree genomes available with contig and scaffold N50s of ∼1.2 Mb and ∼2.1 Mb, respectively. BUSCO analysis, as well as alignment of this reference to other closely related species and to an assembly of tanoak using only third generation reads, suggests that the tanoak draft reference is assembled accurately. Using transcriptome and in silico data, we identified and placed 42,331 genes on the draft reference. In addition, we sequenced a total of 11 unique tanoak trees to better understand the intraspecific diversity. The advanced features of cobarcoded sequencing reads also enabled us to generate haplotype information for each sample with an average N50 of ∼1.6 Mb.

Comparison of the tanoak genome, which has evolved as a species-poor lineage with a limited geographic distribution for a significant period of time, to other related tree genomes (Supplmentary Table 3) showed a similar amount of heterogeneity, which is surprising given the complexity of the evolutionary history of the oaks (*Quercus*), which has involved substantial introgression within a species rich syngameon (Hipp et al, 2020). Interestingly, analysis of R gene composition of tanoak versus other tree genomes showed tanoaks have an overall lower number than many related species and when taking genome size into account, the ratio of R genes to genome size was one of the smallest we measured, particularly among its close relatives in the Fagaceae (Table 3). This low number of R genes may suggest that the tanoak species evolved in a region with relatively few natural parasites or diseases and the absence of introgression for other species with wider distributions and greater exposure to immune system challenges prevented any enrichment or sharing of advantageous alleles. This long-term evolutionary isolation of the tanoak genome may partly explain its rather severe susceptibility to *Phytophthora ramorum*. In addition, a dN/dS comparison of tanoak to pendunculate oak found some positive selection in tanoak for pathogen defense and abiotic stress genes (Plomion et al. 2018), although it is unclear what phenotypic impact this positive selection would have on tanoak.

Using this draft reference, we explored the diversity between different tree samples and identified coding changes that potentially may result in severe alteration of protein function in over 22,000 genes. Comparing this information to SOD resistance in each sample resulted in the discovery of some interesting genes involved in plant immunity and signaling but none reached statistical significance and further studies will be needed to understand individual tree resistance to *Phytophthora ramorum*.

Finally, the unique cobarcoding sequencing method we used enabled the ordering of variants into long haplotypes covering the majority of the genome of each sample. This information resulted in the determination of over 136,000 unique combinations of potentially expressed genes. In all, this reference and the additional samples sequenced have provided a glimpse into the innerworkings of tanoak and provided some sense of the diversity across this species. We hope this reference will be of help to researchers studying tanoak, especially those working to find ways to improve the health and survival of this important species.

## Methods

### DNA isolation

For each sample a single whole leaf was placed in an Oster Pro 1200 blender with 100 ml of lysis buffer (13 mM Tris-HCl (pH 8.3), 140 mM NaCl, 3 mM KCl, 350 mM sucrose, 1 mM EDTA, and 1% Triton X-100) and blended on high for 5 minutes. Lysates were pelleted at 2900 X g for 15 minutes. Supernatants were discarded and the pellet was further isolated using a Nanobind Plant Nuclei Big DNA kit (Circulomics, Baltimore, MD) following the manufacturer’s protocol.

Samples were incubated with Proteinase K for 2 hours, eluted in 100 ul of elution buffer, and quantified using a Nanodrop 1000 spectrophotometer (ThermoFisher, Waltham, MA). To enrich for longer DNA molecules 6 samples were further processed using a Short Read Eliminator XL kit (Circulomics, Baltimore, MD) prior to making stLFR libraries. This long fragment enriched DNA was also used for the Minion sequencing (Oxford Nanopore Technologies, Oxford, UK).

### Cobarcoded read libraries

Cobarcoded read libraries were generated using an MGIEasy stLFR Library Prep kit (MGI, Shenzhen, China) following the manufacturer’s protocol using 1 ng of input DNA. stLFR libraries were analyzed on a DNBSEQ-G400 (MGI, Shenzhen, China) DNA sequencer using pair-end 100 base reads and a 42 base barcode read. stLFR fq files were processed using the barcode split tool (GitHub; https://github.com/stLFR/stLFR_read_demux) (Wang et al. 2019) to deconvolute barcodes.

### Nanopore libraries

Minion libraries were prepared using the Genomic DNA by Ligation kit (Oxford Nanopore, Oxford, UK) following the recommended protocol. Briefly, the isolated DNA was first repaired and end-preprepared by the NEBNext FFPE DNA Repair mix (New England Biolabs, Ipswich, MA) and NEBNext Ultra II End repair/dA-tailing Module (New England Biolabs, Ipswich, MA) following the manufacturer’s protocol. The reaction was purified using a 1X volume of Ampure XP beads (Beckman Coulter, A63882) following the manufacturer’s protocol. The product was then ligated with the Adapter Mix and purified with an optimized protocol provided by the Genomic DNA by Ligation kit. After purification the library was ready for sequencing.

Minion sequencing was carried out following the manufacturer’s suggested protocol. The priming buffer mix was first prepared in accordance with the protocol and then loaded onto a R9.4.1 flow cell. The final sequencing library was prepared by mixing 50 fmol of purified library with the sequencing buffer and the loading beads. The loaded flow cell was then mounted onto a MinION Mk 1B device (Oxford Nanopore, Oxford, UK) and sequenced with MinKNOW v19.10 for approximately 24 hours. FAST5 files were analyzed with Guppy and configuration file dna_r9.4.1_450bps_fast.cfg.

### Genome assembly

Cobarcoded sequencing reads from stLFR data were assembled using a modified version of Supernova (10X Genomics, Pleasanton, CA) that allows for greater than 4 million unique barcodes. The six Supernova assemblies generated from DNA enriched for long fragments were used to build a single genome assembly for tanoak by using contigs from NL.2.XL, SM.52.81.XL (clone 2), LP.22.48.XL, SM.52.42.XL, and SM.54.37.XL to fill gaps in the SM.74.45.XL (clone 1) assembly. This was performed using TGS-gapfiller using standard settings. To further improve the pangenome assembly TGS-gapfiller (Xu et al. 2020) was used with 11.8, 9.6, and 6 Gb of Minion generated reads from SM.54.37.XL, SM.52.82.XL (clone 2), and NL.2.XL, respectively. Finally, SM.74.45.XL (clone 1) cobarcoded reads were aligned to the genome assembly with bwa and the genome was polished using pilon.

We observed 2 possible mis-assemblies on the dotplot (ONT contig_83 aligned to draft genome on both 262_pilon and 656_pilon; contig_513 aligned to 360_pilon and 819_pilon). We aligned the ONT reads to both the ONT assembly and the draft genome but found no confident evidence to indicate which assembly is the correct one. Future investigation with improved sequencing techniques will be necessary to disambiguate this region.

After performing joint calling of all 19 sequencing libraries (see “Variant calling and Phasing” below), we discovered 58,314 homozygous alternative allele variants shared by all 19 libraries. This suggested that these alternative alleles should in fact be the reference allele. To correct this, we replaced all 58,134 positions in the reference with the alternative allele and created a new v2 reference. For most analyses this change was immaterial and v1 was continued to be used, but for all variant calling applications v2 was used.

### Genome analysis

Genome completeness and contiguity was analyzed with BUSCO version 5.2.2 (Manni et al. 2021) using standard options with embryophyta_odb10 data set. N50 statistics and other genome metrics were generated using QUAST version 5.0.2 using default settings.

The draft genome was aligned to *Q. robur* and *Q. rubra* genomes with minimap2 v.2.16-r922 (Li 2018). The alignment results were then visualized with pafCoordsDotPlotly.R from dotPlotly (https://github.com/tpoorten/dotPlotly).

### Genome annotation

Protein-coding gene annotation was performed using MAKER v 3.01.04 (Campbell et al. 2014b; Cantarel et al. 2008). A *de novo* transcriptome assembly, protein sequences from two related oak species (Plomion et al. 2018), a tanoak repetitive elements library, and a tanoak gene prediction model were used as gene evidence for the initial round of gene prediction with default parameters. The *de novo* transcriptome was assembled by Trinity v2.8.5 (Grabherr et al. 2011) with input mRNA from a SRA study SRP157197 (Kasuga et al. 2021) of 45 samples. Protein sequences came from *Q. robur* (English Oak) and *Q. rubra* (Northern Red Oak). The repeat library was established using RepeatMasker v4.0.7 (A.F.A. Smit, R. Hubley & P. Green RepeatMasker at http://repeatmasker.org) following the MAKERP pipeline method (Campbell et al. 2014a). The gene prediction model was created using the BUSCO v4.1.4 (Manni et al. 2021) pipeline with AUGUSTUS (Stanke and Waack 2003) in genome mode with the lineage eukaryota_odb10. The resulting gene models from MAKER were then used to train SNAP (Korf 2004) and create an HMM file. MAKER was then used for a second round of gene prediction using the previously mentioned gene evidence along with the first round MAKER annotations and SNAP HMM file. The resulting gene models were then filtered to keep annotations with an AED <= 0.5.

The MAKER generated proteins were compared against UniProt/SwissProt database (UniProt 2021) with BLASTP (BLAST v2.13.0+) to get homology-based annotation. Interproscan v5.59.91.0 was used to identify protein domains and predicted GO terms. A total of 51,233 protein-coding genes were identified.

### Variant calling and Phasing

The Genome Analysis ToolKit (GATK) (v.4.1.2.0) was used for variant calling. For each sample, the HaplotypeCaller function was used to call GVCF files. After combining all GVCF files, the GenotypeGVCFs function was used to joint genotype variants. Variants were hard-filtered to keep >=15x coverage across all samples. Low quality variants were removed with QD<2.0|| MQ<26.0|| FS>100.0|| SOR>5.0|| MQRankSum< -7.5|| ReadPosRankSum< -8.0 (parameters adopted from (Hu et al. 2022)). The resulting high-quality variants were phased with Hapcut2 v.1.3 (Bansal 2023) for each sample. Due to high diversity and variant calling errors, two haplotypes are considered the same if they share greater than 90% similarity of SNP calls.

### Polymorphism Analyses

Scripts for the SNP association tests, SnpEff annotation, *Arabidopsis* annotation, PCA, and dN/dS analyses described below are available at https://github.com/MaloofLab/Cai-TanOak-2023. SnpEff v5.1d (Cingolani et al. 2012) was used to predict the possible consequences of each SNP on protein coding genes. To determine the closest *Arabidopsis* homolog for each tanoak gene, blastp (Altschul et al. 1990; Altschul et al. 1997) was used to blast the tanoak proteome against Arabidopsis TAIR10 protein sequences (downloaded from ftp://ftp.arabidopsis.org/home/tair/Sequences/blast_datasets/TAIR10_blastsets/TAIR10_pep_20110103_representative_gene_model_updated.)

## Data Availability

Co-barcoded and ONT sequencing data generated for this study have been deposited in the SRA under BioProject PRJNA944640. The Notholithocarpus densiflorus draft assembly has been deposited at DDBJ/ENA/GenBank under the accession JARYZH000000000. For most analyses version described in this paper is JARYZH010000000. For variant calling an updated version JARYZH010000000@@@ was generated that corrected alternative homozygous variants found in all samples to be the reference bases. Scripts for the SNP association tests, SnpEff annotation, *Arabidopsis* annotation, PCA, and dN/dS analyses are available at https://github.com/MaloofLab/Cai-TanOak-2023.

## Acknowledgements

USDA-NIFA award CA-D-PLB-2795-H was used to support J.N.M.’s work on this project.

